# N-Zinclopedia: A database of predicted transition metal-binding sites and metalloenzymes

**DOI:** 10.64898/2026.10.18.759936

**Authors:** Gilad Mechtinger, Gabriel Axel, Larson Marcellino, Alexander M. Bowman, Joanna Slusky, Nir Ben-Tal

## Abstract

Metal ions are among the most common enzymatic cofactors, yet metal utilization remains poorly mapped. Here, we present N-Zinclopedia, an atlas of predicted transition metal-binding sites and candidate metalloenzymes across protein fold space. N-Zinclopedia catalogs protein sites compatible with transition metal coordination, identified using ZincSight screening of protein structure model databases. We focused on previously uncharacterized domains and proteins, screening for transition metal-binding sites within over a million of these, spanning both poorly annotated and canonical folds. We also classified the predicted sites, using MAHOMES-II, as candidate catalytic or non-catalytic metal-binding sites. Experimental characterization of a previously unknown zinc binding site confirms that ZincSight can determine previously un-annotated zinc binding. N-Zinclopedia significantly expands the known landscape of metal utilization, providing numerous predictions that await experimental examination. N-Zinclopedia is available at: https://bentallab.github.io/nzinclopedia.

## Introduction

Metalloproteins are estimated to account for 30-40% of proteins in proteomes, and metalloenzymes for roughly 40% of all enzymes (1, 2). Despite this prevalence, metal utilization remains incompletely mapped across protein space (3–7). Curated resources, including MetalPDB (8), and InterMetalDB (9), ZincBind (10), MeDBA (11), MESPEUS (12) and BioLip2 (13), catalog experimentally observed metal-binding sites, while state-of-the-art predictors such as Metal3D (14) and PinMyMetal (15) enable per-protein identification of candidate sites. Nevertheless, an atlas-scale resource that systematically maps predicted metal-binding sites across large and diverse protein collections is still lacking. Compounding this, UniProtKB reports residue-level metal-binding annotations for ∼17% of curated proteins but only ∼3% of unreviewed entries, underscoring the extent of missing site-level annotations at the proteome scale (16).

As the volume of known protein/metal ion complexes has rapidly increased in recent years, systematic high-throughput mapping of transition metal-binding sites remains limited; possibly because of the inherent trade-off between prediction accuracy and the computational scalability required for proteome-scale screening (4, 5, 7, 16, 17). Here, we address this gap using our recently introduced transition metal-binding site predictor, ZincSight (18). We applied it at scale across datasets spanning both novel and canonical folds. Specifically, we analyzed more than one million structures, including 711,705 representative proteins from AlphaFold Database (AFDB) Dark Clusters (19–23); 615,571 representative domains comprising three widespread fold topologies: 570,853 Rossmann-like domains, 32,271 TIM-barrel domains, and 12,447 seven-bladed β-propeller, methylamine dehydrogenase chain H-like domains; and 7,427 representative domains from novel-fold clusters in The Encyclopedia of Domains (TED) (24).

Following our high-throughput ZincSight predictions, to support functional interpretation of predicted sites, we incorporated Metal Activity Heuristic of Metalloprotein and Enzymatic Sites (MAHOMES-II) (25, 26), which classifies metal-binding environments as catalytic or non-catalytic using local structural and physicochemical descriptors. Together, ZincSight and MAHOMES-II enable systematic identification and classification of candidate metalloenzyme active-sites, as well as putative structural or regulatory metal-binding motifs. To provide additional support for selected high-confidence candidates, we further performed targeted AlphaFold3 (27) zinc-complex modeling on a subset of predicted metalloenzymatic sites.

## Results

### Detection of metalloenzymatic TED novel domains

To explore candidate metalloenzymatic sites from a fold-novelty perspective we analyzed TED novel-fold domain representatives. TED partitions AFDB into nearly 365 million domains grouped into approximately 120 million clusters. Of these, about 78 million clusters received a CATH-based assignment, whereas ∼41 million remained unlabeled. Because not all CATH-unlabeled domains represent true novel folds, TED applied additional filtering and structural-similarity searches, with exhaustive TM-align as a final sensitive step, yielding 7,427 putative novel-domain clusters. We focused on these domains because they occupy structurally distinct regions of protein space and therefore provide a compelling setting for identifying candidate metal-binding sites not readily attributable to known metalloprotein families.

Analysis of the 7,427 TED novel-fold representative domains showed that zinc-compatible coordination geometries recur broadly across previously uncharacterized folds. Using the dual-gate thresholds defined in Methods, 1,359 sites passed the baseline criterion on the unrefined structures, of which 852 also passed the stricter histidine-rotamer-refined criterion, yielding a high-confidence set distributed across 615 unique structures.

We then used MAHOMES-II to separate sites likely to be metalloenzymatic from those serving structural or regulatory roles. At a catalytic-probability threshold of p_(MAHOMES)_ ≥ 0.5, seventy-one sites were classified as putatively catalytic. Visual inspection showed that these sites consistently occupy pocket-like cavities, ranging from relatively solvent-exposed clefts to deeply buried environments, rather than broad, featureless surface patches, consistent with canonical enzyme active-site localization.

We next assessed whether these candidates represent genuinely unannotated metalloenzymes or rediscoveries of known function. Foldseek (28) structural comparison eliminated 35 sites whose closest structural neighbors carried existing metalloenzymatic annotations. The remaining Foldseek-filtered candidates were then examined by targeted AlphaFold3 zinc-complex modeling; 12 were excluded because the modeled zinc ion fell below the local confidence threshold of pLDDT ≥ 90. ConSurf (29–31) analysis showed that predicted coordinating residues were consistently highly evolutionarily conserved, consistent with their functional importance (Table 1).

**Table 1.** Predicted transition metal-binding sites and inferred catalytic propensity for selected UniProt proteins. For each UniProt ID, ZincSight reports the predicted probability (%) that the listed residue set coordinates a zinc ion (or other transition metal), shown both with and without histidine imidazole rotamer sampling (“With His rotamer” and “Without His rotamer”). Binding Residues lists the predicted coordinating residues (chain and residue index), and Binding Residue Types indicates their amino-acid identities in one-letter code. MAHOMES-II Catalytic Probability (%) reports the predicted likelihood that the site is catalytic. Multiple rows for the same UniProt ID correspond to distinct predicted sites within that protein. Secondary sub-threshold sites are shown in grey; these sites are nevertheless supported by high-confidence AlphaFold3 models (pLDDT > 90). For each predicted binding site, the table also lists the ConSurf conservation grades of the coordinating residues in a 1 to 9 scale, where 9 is the most highly conserved. The table further reports the number of homologous sequences recovered by the AlphaFold2-integrated homologue-search workflow, including JackHMMER (32, 33) and HHblits (34).

| Uniprot ID | ZincSight without His rotamer probability (%) | ZincSight with His rotamer probability (%) | Binding residues | Binding residues types | ConSurf conservation score | MAHOMES II catalytic probability (%) | Number of homologues |
| --- | --- | --- | --- | --- | --- | --- | --- |
| A0A0M5K0Y8 | 45.91 | 69.23 | A_60;<br>A_58;<br>A_48 | {D, H, H} | 9;9;9 | 100 | 1054 |
| A0A0M5K0Y8 | 26.06 | 37.78 | A_143;<br>A_62;<br>A_112 | {E, H, H} | 9;9;9 | 100 | 1054 |
| A0A1I3EZC5 | 41.82 | 43.25 | A_94;<br>A_116;<br>A_114 | {E, H, H} | 9;9;9 | 100 | 2268 |
| A0A1I3EZC5 | 0.7 | 10 | A_94;<br>A_66;<br>A_52 | {E, H, H} | 9;9;9 | 100 | 2268 |
| A0A1N7Q4L0 | 55.08 | 87.92 | A_124;<br>A_95;<br>A_96 | {D, H, H} | 9;9;8 | 99.93 | 2222 |
| A0A2H5WW51 | 83.26 | 83.26 | A_372;<br>A_306;<br>A_302 | {E, H, H} | 9;9;9 | 100 | 48 |
| A0A2J7VJ59 | 55.89 | 60.86 | A_54;<br>A_43;<br>A_38 | {E, H, H} | 9;9;9 | 100 | 23 |
| A0A3C0G6M2 | 79.25 | 86.39 | A_152;<br>A_131;<br>A_126 | {E, H, H} | 9;9;9 | 100 | 169 |
| A0A4Z1HKJ8 | 63.73 | 63.73 | A_517;<br>A_825;<br>A_515 | {D, H, E} | 9;9;9 | 100 | 1797 |
| A0A536G3B6 | 69.86 | 69.86 | A_120;<br>A_65;<br>A_61 | {D, H, H} | 9;9;8 | 99.8 | 7316 |
| A0A545UDA7 | 40.81 | 87.05 | A_185;<br>A_183;<br>A_176 | {H, H, H} | 9;9;9 | 99.87 | 812 |
| A0A7C2D0E3 | 90.54 | 90.54 | A_99;<br>A_65;<br>A_222 | {D, H, E} | 9;9;9 | 99.72 | 351 |
| A0A7S0R8P6 | 48.34 | 48.34 | A_135;<br>A_133;<br>A_123 | {D, H, H} | 9;9;9 | 100 | 474 |
| A0A7S0R8P6 | 0.54 | 23.83 | A_137;<br>A_197;<br>A_229 | {E, H, H} | 9;9;9 | 100 | 474 |
| A0A7V1D5I9 | 50 | 50 | A_408;<br>A_380;<br>A_389 | {D, H, E} | 9;9;9 | 100 | 2967 |
| A0A7Y5NWC9 | 35.66 | 56.86 | A_215;<br>A_259;<br>A_261 | {E, H, H} | 9;8;9 | 98.01 | 376 |
| A0A7Y6ZTD4 | 77.22 | 87.13 | A_329;<br>A_327;<br>A_306;<br>A_350 | {H, H, H,<br>E} | 9;9;9;9 | 99.94 | 537 |
| A0A7Y7D8I8 | 64.56 | 90.1 | A_269;<br>A_107;<br>A_103 | {E, H, H} | 9;9;9 | 100 | 292 |
| A0A840Y722 | 82.2 | 88.14 | A_104;<br>A_52;<br>A_48 | {E, H, H} | 9;9;9 | 100 | 109 |
| E4YIH6 | 59.52 | 69.64 | A_135;<br>A_133;<br>A_126 | {H, H, H} | 9;9;9 | 100 | 828 |
| L8WQG2 | 45.96 | 55.3 | A_402;<br>A_340;<br>A_338 | {C, H, H} | 9;9;9 | 99.89 | 206 |
| N9UBA4 | 52.67 | 83 | A_177;<br>A_142;<br>A_181 | {H, H, H} | 9;9;9 | 99.87 | 21 |
| W9DNB8 | 81.08 | 81.08 | A_109;<br>A_53;<br>A_49 | {E, H, H} | 9;9;9 | 100 | 33 |
| A0A0H4B904 | 26.71 | 57.9 | A_337;<br>A_357;<br>A_353 | {H, H, H} | 9;9;9 | 99.81 | 841 |
| A0A497LQG5 | 28.31 | 77.2 | A_35;<br>A_33;<br>A_4 | {D, H, H} | 9;9;9 | 99.89 | 60 |
| A8FZD8 | 29.94 | 79.67 | A_111;<br>A_74;<br>A_109 | {H, H, H} | 8;9;9 | 99.98 | 42 |
| A0A7S3TQB6 | 24.55 | 43.41 | A_448;<br>A_465;<br>A_450 | {H, H, H} | 9;9;8 | 84.38 | 217 |

This multi-stage filtering yielded 24 high-confidence candidate metalloenzymatic sites, each satisfying four independent criteria: local geometric compatibility with zinc coordination, physicochemical features predictive of catalysis, absence of an obvious structural explanation based on annotated metalloenzymes, and complementary support from AlphaFold3 complex modeling and evolutionary conservation (Figure 1).

**Figure 1.**
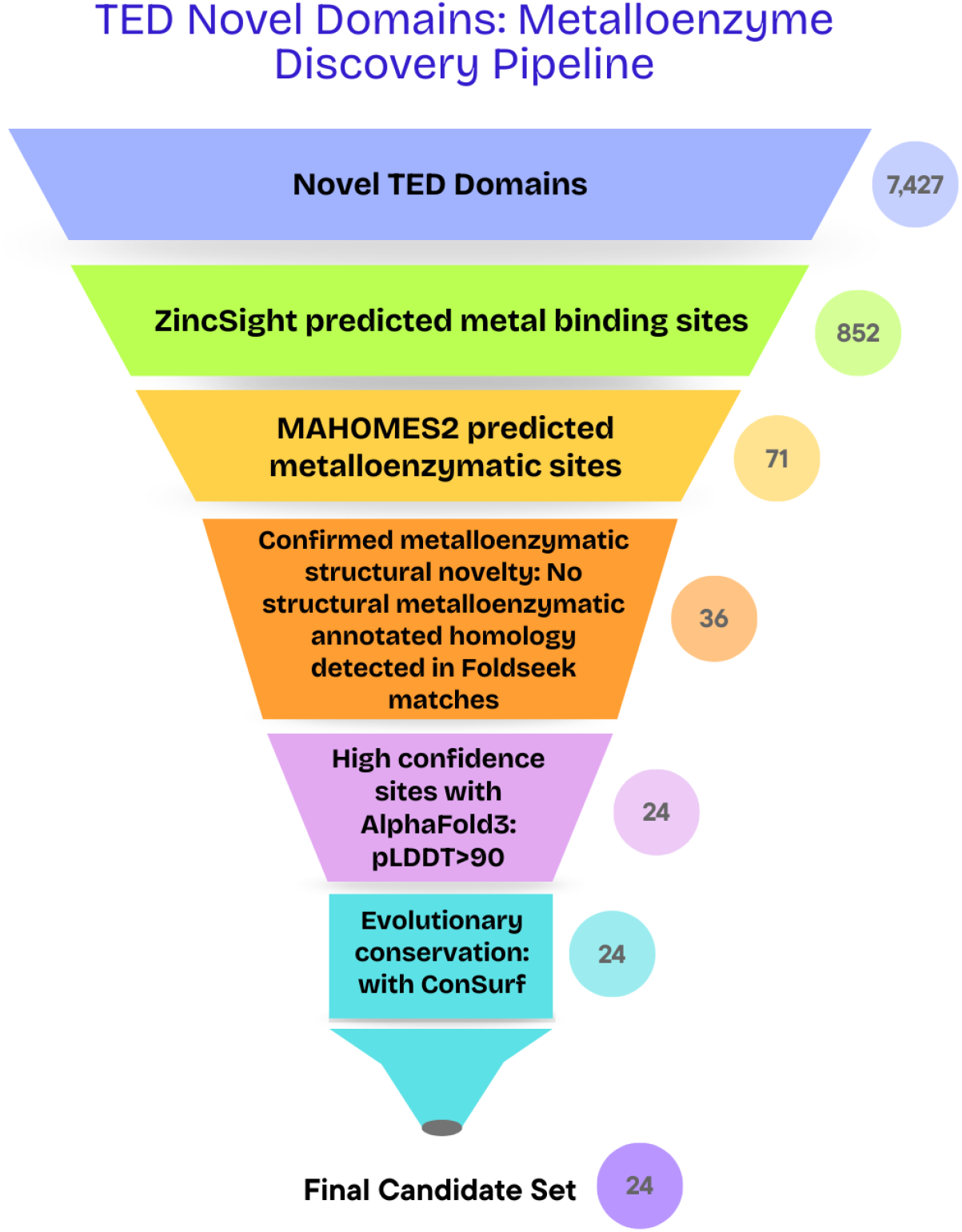
TED novel-domain metalloenzyme discovery pipeline. From 7,427 TED novel domains, ZincSight identified 1,359 candidate metal-binding sites, of which 852 passed the stricter rotamer-refined criterion. MAHOMES-II classified 71 as putatively catalytic (p_(MAHOMES)_ ≥ 0.5), and Foldseek filtering retained 36 sites lacking annotated metalloenzymatic structural homologues. Targeted AlphaFold3 zinc-complex modeling of the Foldseek-filtered candidates (zinc ion pLDDT ≥ 90), together with ConSurf conservation analysis, yielded a final set of 24 high-confidence candidate metalloenzymatic sites.

Importantly, these candidates were not confined to rare or isolated proteins. Eighteen of the 24 metal binding sites were predicted in representatives of protein families with more than 100 homologues. The remaining predicted sites occurred in smaller or less well-sampled lineages. This distribution implies that unannotated metalloenzymatic potential is not idiosyncratic but instead extends across evolutionarily broad families that have so far escaped metal-related functional characterization (Table 1).

#### Stoichiometry sensitivity in targeted AlphaFold3 follow-up models

AlphaFold3 was not used in the initial database-scale screening workflow, but only as a targeted follow-up step for a selected subset of candidates that had already passed ZincSight screening, MAHOMES-II catalytic classification, and Foldseek-based novelty filtering. Within this subset, confidence at the modeled zinc-ion position was sensitive in some cases to the assumed metal-ion stoichiometry.

Three proteins, A0A0M5K0Y8, A0A7S0R8P6, and A0A1I3EZC5, initially yielded zinc-ion pLDDT values below 90, but re-modeling with a second zinc ion restored high local confidence in each case, suggesting possible binuclear configurations. Consistently, ZincSight identified a second sub-threshold candidate site in each protein, reinforcing the interpretation that these structures may bind two transition metals.

The modeled pocket geometries differed between cases. For the TTHB210-like proteins A0A0M5K0Y8 and A0A7S0R8P6, both candidate sites remained detectable in metal-free AlphaFold3 models, indicating that the local pocket geometry is intrinsically stable and does not depend on prior metal occupancy. A0A1I3EZC5 behaved differently: the secondary site was recovered by ZincSight only in the one-zinc model, raising the possibility that initial metal binding induces a local conformational rearrangement that favors formation or stabilization of the second site. For transparency, these secondary sub-threshold sites are listed in grey in Table 1.

### Representative case studies from TED novel-fold candidates

A0A1I3EZC5 (Figure 2A), which maps to DUF1264 (PF06884), is among the more structurally compelling candidates in our set and is associated with at least 2,268 homologues (Table 1). The predicted site is binuclear and it resides within a deep, enclosed pocket - a geometry more consistent with small-molecule substrate accommodation than with the extended, shallow clefts typical of enzymes acting on macromolecular substrates. Pocket accessibility visual analysis reinforces this interpretation: the surface computed with the default 1.4 Å solvent radius indicates an occluded entrance in the static model, whereas a slab-clipped view reveals that the predicted site lies within a deep, buried pocket. The DUF1264 family, spans bacterial and eukaryotic organisms. Its members are typically ∼200 residues long. Some of these carry lipoprotein annotations, suggesting possible membrane anchoring in some lineages. Notably, STRING (35) reports predicted functional associations between A0A1I3EZC5 and the glyoxalase-pathway enzymes, GloB (∼0.845) and glyoxalase I (∼0.607), involved in glutathione-dependent methylglyoxal detoxification (36). Taken together, the pocket geometry and network context are consistent with a role in carbonyl-stress metabolism, though this remains a hypothesis to be tested experimentally.

**Figure 2.**
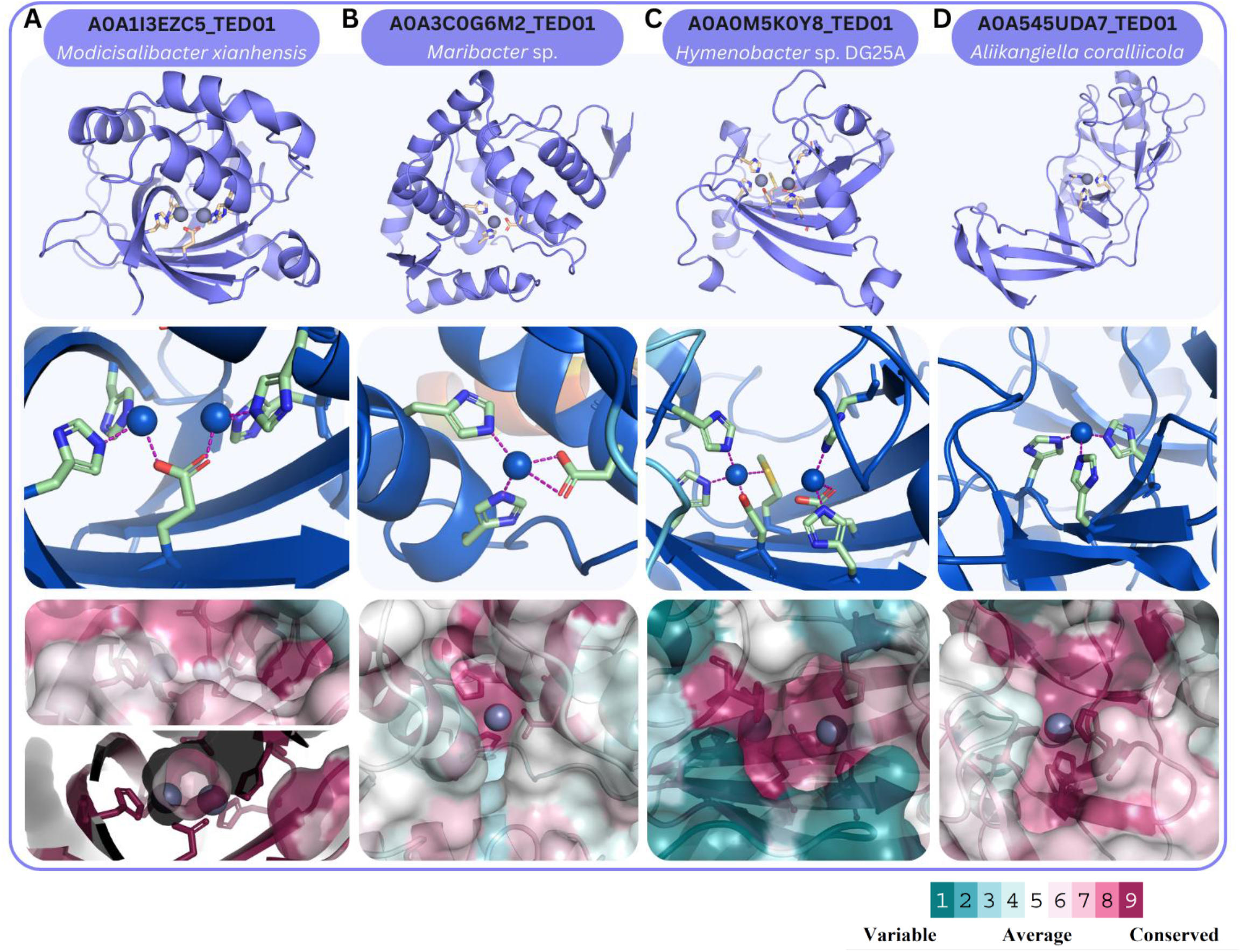
Targeted AlphaFold3 zinc-complex models for selected TED-domain candidate metalloenzymes: **(A) A0A1I3EZC5, (B) A0A3C0G6M2, (C) A0A0M5K0Y8, and (D) A0A545UDA7. Top:** overall protein folds with predicted zinc ion positions shown as grey spheres; zinc-coordinating residues are shown as wheat-colored sticks. **Middle:** zoom-in of the predicted sites with coordinating residues shown as sticks; dashed lines denote zinc–donor coordination interactions. Models are colored by the AlphaFold pLDDT scale; the zinc-ion spheres are colored on the same scale to reflect local confidence at the predicted ion positions. The zinc-ion pLDDT values were 98.73 and 98.49 in panel A, 94.61 in panel B, 93.37 and 94.98 in panel C, and 95.81 in panel D. **Bottom**: surface representations colored according to the ConSurf conservation scale (color bar shown), with zinc ions (grey) positioned within putative binding pockets rather than on exposed surface regions; surfaces were computed in PyMOL using the default 1.4 Å solvent radius. For A0A1I3EZC5, an additional slab-clipped panel reveals that the predicted site lies within a buried pocket.

A0A3C0G6M2 (Figure 2B), represents a mononuclear candidate for which structural analysis alone does not immediately suggest a substrate class. As a preliminary functional pointer, DeepFRI (37) predicts purine or nucleotide binding (score 0.74) and broad metabolic process involvement (up to 0.77), while EZPred (38) tentatively favors a transferase classification over hydrolase or lyase. These signals are not definitive and should not be over-interpreted, but their directional consistency, pointing toward nucleotide-associated chemistry, provides a reasonable starting hypothesis for biochemical follow-up, particularly in designing substrate screens.

Two additional candidates, A0A0M5K0Y8 (Figure 2C) and A0A7S0R8P6, are both annotated by Pfam (39) as members of the TTHB210-like domain family. The only experimentally solved structure in this family, 3VG8, contains no metal ions (40), leaving metal dependence an open question. Our structure-based predictions indicate that both proteins harbor metalloenzymatic candidate sites, raising the possibility that metal-dependent activity exists within this family and has so far gone undetected.

Finally, A0A545UDA7 (Figure 2D) is predicted to contain a zinc-binding site coordinated by three histidine residues. Its primary interest lies at the family level: Pfam assigns it to the YHYH domain family (PF14240), which currently lacks metal-binding and enzymatic characterization, making it a well-motivated blank-slate target for experimental follow-up.

### Detection of uncharacterized metalloenzymatic sites in canonical folds

While TED novel domains represent unprecedented protein architectures, structural novelty does not exhaust the landscape of uncharacterized metal utilization. Even within well-established enzyme folds, metal dependence remains incompletely annotated. We therefore extended our standardized screening pipeline to canonical enzyme fold classes, identifying candidate metalloenzymatic sites within established folds that remain functionally underexplored.

To enrich previously uncharacterized metalloenzymatic domains, we focused on AlphaFold2 TED domains clustered at the Foldseek-derived T-level (topology) rather than the H-level (homology). While H-level clusters typically correspond to well-characterized protein families, restricting our analysis to T-level topologies increases the likelihood of capturing remote domains that remain functionally unannotated. Within three major topology groups: TIM-barrel (CATH 3.20.20), 7-bladed β-propeller/methylamine dehydrogenase chain H-like (CATH 2.130.10) and Rossmann-like fold (CATH 3.40.50), we screened for catalytic metal-binding candidates using ZincSight followed by MAHOMES-II. We then assessed these shortlisted candidates using the same Foldseek-based structural control and targeted AlphaFold3 zinc-complex modeling workflow applied in the TED novel-domain analysis. From these results, we selected three illustrative examples, one from each fold: A0A193LBS4-TED01 (TIM-barrel), A0A847GYA2 (β-propeller), and A0A4U8TDW7 (Rossmann-like), as highlighted in Figure 3.

**Figure 3.**
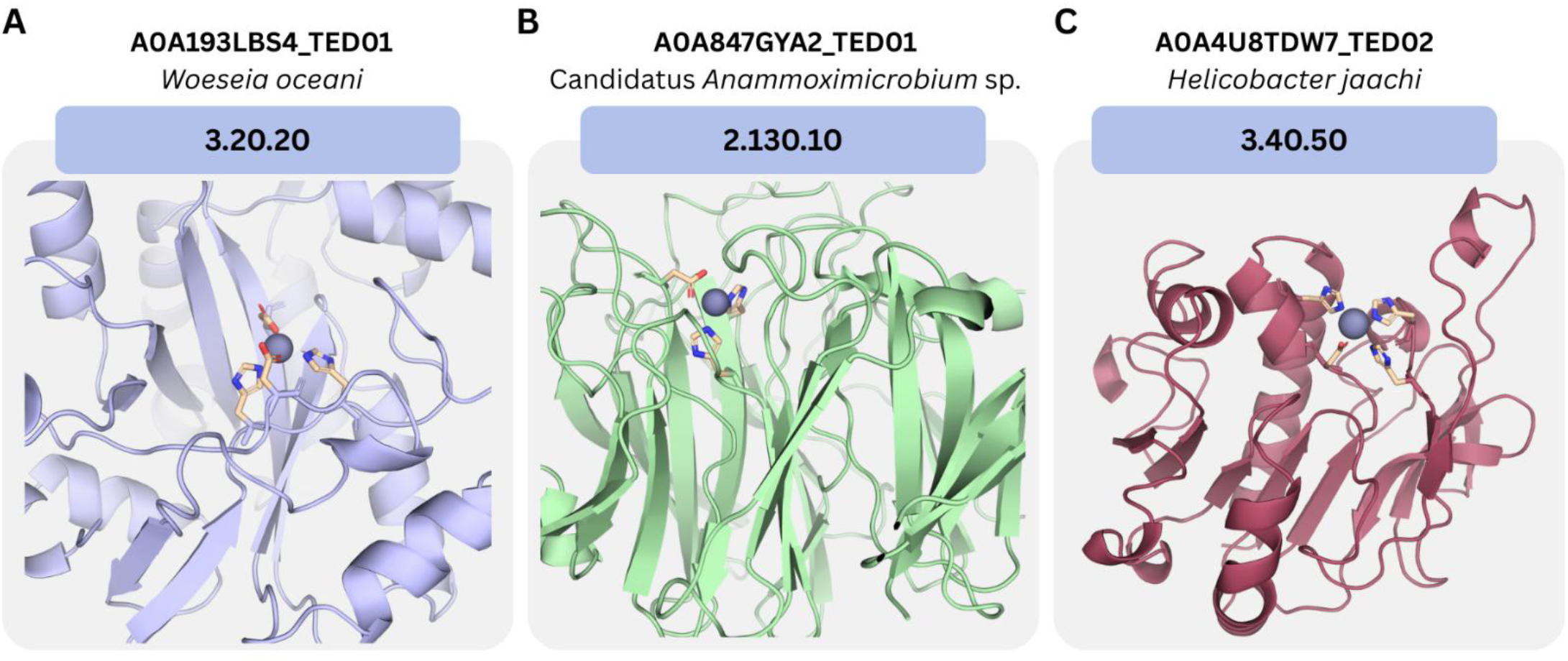
Targeted AlphaFold3 zinc-complex models of selected metalloenzymatic candidate domains representing three canonical topology groups. (A) TIM-barrel (CATH 3.20.20; A0A193LBS4_TED01), (B) 7-bladed β-propeller, methylamine dehydrogenase chain H-like (CATH 2.130.10; A0A847GYA2_TED01), and (C) Rossmann-like (CATH 3.40.50; A0A4U8TDW7_TED02), each modeled in complex with a zinc ion. Although each domain matches a known global fold, Foldseek retrieves structurally remote neighbors that explain the overall topology but do not support the predicted metal-binding architecture or a related functional annotation. The modeled zinc ions have high confidence (pLDDT > 90); zinc pLDDT values are 97.77, 91.50, and 97.71 for panels A-C, respectively. Proteins are shown as cartoons in a uniform color for clarity, and zinc-coordinating residues are shown as wheat-colored sticks.

Across all three topology groups, the novelty signal follows a consistent pattern: Foldseek retrieves high-confidence structural neighbors that explain the global fold, yet these matches do not recapitulate the predicted metal-binding geometry or carry related metalloenzymatic annotations. This divergence suggests the existence of previously unrecognized metalloenzymatic lineages or active-site architectures embedded within established fold scaffolds. Notably, Rossmann-like folds are less frequently associated with transition-metal-dependent catalysis than TIM barrels or β-propellers, making the Rossmann-like candidate particularly compelling. Even within these well-established fold classes, some protein families remain poorly characterized. For example, A0A1Y5S9R6, of the TIM-barrel fold, belongs to the poorly characterized DUF3604 family, for which metal binding or metalloenzymatic activity has not been reported. Together, these cases illustrate that potentially unrecognized metal-dependent functions can occur even within otherwise well-established fold classes.

### Screening unannotated proteins from AFDB dark clusters

TED novel-domain and canonical-fold analyses broaden structural coverage but remain domain-resolved. Putative novel folds are restricted to high- or medium-consensus, CATH-unlabelled domains that pass stringent quality and similarity filters, whereas canonical-fold analyses cover established fold classes.

Consequently, many AlphaFold DB proteins remain sparsely annotated and fall outside both analysis sets. To investigate this complementary protein-level space, we screened representative proteins from AFDB dark clusters. Starting from 2.30 million non-singleton AFDB structural clusters, we excluded clusters whose representative matched a known PDB structure or Pfam family, as well as clusters containing any member with a Pfam or TIGRFAM annotation in UniProtKB/TrEMBL or UniProtKB/Swiss-Prot. This filtering yielded 711,705 dark clusters, each represented by a single protein.

As representative examples, we highlight three candidates predicted to be metalloenzymatic: A0A643J265, A0A835KIB7, and A0A4U0H974 (Figure 4). These cases are notable because they fall outside both TED analysis sets: A0A643J265 contains a predicted site whose binding residues lie partly within a TED domain, whereas A0A835KIB7 and A0A4U0H974 harbor predicted sites within TED domains that are neither designated as novel folds nor assigned to any CATH topology. We then assessed novelty of these shortlisted candidates using the Foldseek-based searches and AlphaFold3 zinc-complex modeling workflow.

**Figure 4.**
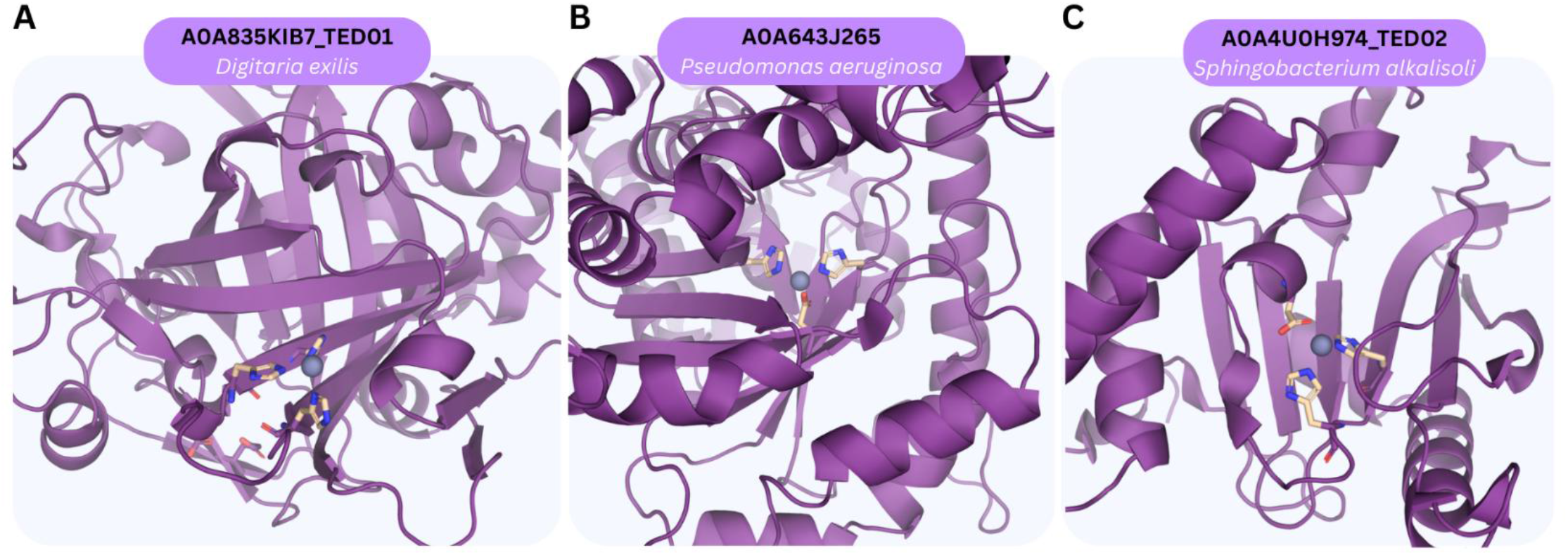
Representative examples of ZincSight-predicted metal-binding sites in AFDB dark-clusters proteins. These cases illustrate how the screening workflow extends beyond TED-defined and canonical domain sets into annotation-poor protein space. Shown are (A) A0A835KIB7_TED01, (B) A0A643J265 and (C) A0A4U0H974. Zinc-coordinating residues are shown as wheat-colored sticks.

### Experimental characterization

To determine the veracity of predictions of previously uncharacterized zinc sites, we experimentally characterized a candidate protein and found that it bound zinc although at a lower zinc-to-protein ratio than predicted. Proteins A0A9X3MNF5 and A0A3W0S055, homologues of A0A3C0G6M2 and A0A1I3EZC5, respectively, were selected for experimental validation because of their high ZincSight scores and practical considerations for expression. A0A9X3MNF5 was selected for its shorter sequence and higher mean pLDDT in its full-length AlphaFold model compared with A0A3C0G6M2. A0A3W0S055 was selected because its source organism, Salmonella enterica, is closely related to the E. coli expression host. Because SignalP 6.0 (41) predicted an N-terminal signal peptide in A0A3W0S055, the protein was expressed without its first 17 amino acids.

Expression and purification were attempted for both proteins. Purification methods focused on optimizing conditions for ammonium sulfate precipitation followed by dialysis, Q-column chromatography, and size-exclusion chromatography (see *Methods*). A0A9X3MNF5 was expressed but consistently aggregated. A0A3W0S055 was successfully expressed and purified (Figure 5A and B). The mass of A0A3W0S055 was measured by MALDI mass spectrometry. The expected molecular weight of the protein is 27,181.60 Da. The dominant peaks at 26,945 and 27,094 Da may reflect cleavage of terminal amino acids (Figure 5C).

**Figure 5.**
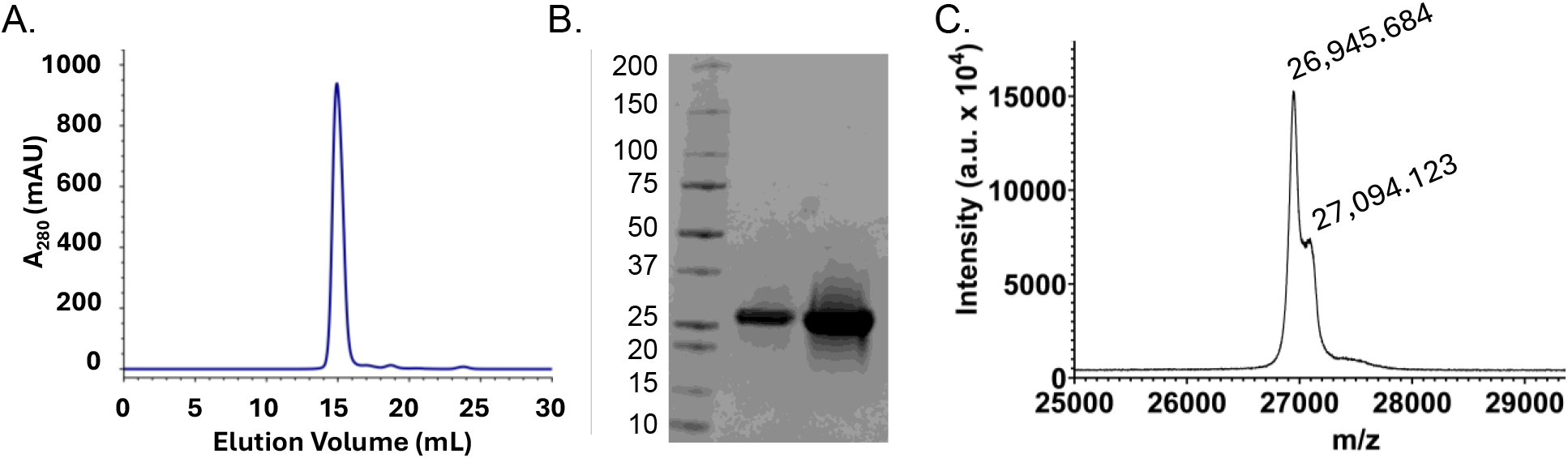
Purification of A0A3W0S055. **A.** Size-exclusion chromatogram shows a unimodal peak at an elution volume consistent with pure protein B. Protein purity was confirmed by sodium dodecyl sulfate polyacrylamide gel electrophoresis (SDS-PAGE) with 2 μg and 10 μg of protein C. MALDI mass spectrometry confirmed the identity of the protein.

Metal content of the protein was analyzed by Inductively Coupled Plasma Optical Emission Spectroscopy (ICP-OES) before and after soaking in a ZnCl2 solution (see Methods). A semiquantitative ICP-OES analysis of biological transition metals revealed that Zn was the predominant transition metal bound to the protein. Quantitative ICP-OES revealed the zinc-to-protein molar ratio was 0.5 moles of zinc per mole of (monomeric) A0A3W0S055. After soaking in a solution of ZnCl2 and then desalting the protein, we found the zinc-to-protein molar ratio was 0.91 moles of zinc per mole of (monomeric) A0A3W0S055. All metal quantification used a control of the buffer that had also passed through the last step of purification—desalt column for soaked protein, size exclusion column for unsoaked protein—for baseline correction. A0A3W0S055 was predicted by ZincSight to bind two zinc ions per monomer of protein.

### The N-Zinclopedia database

N-Zinclopedia (https://bentallab.github.io/nzinclopedia/) is an open-access database of predicted transition metal-binding sites across protein fold space, integrating ZincSight site predictions with MAHOMES-II catalytic classifications (Figure 6A). Users can explore candidate metal-binding sites at residue-level detail through an embedded Mol* viewer (Figure 6B) (42). The database includes representative domains from the TED novel-fold database, and from three canonical T-level topology groups (TIM barrel, β-propeller, and Rossmann-like). It also includes representative proteins from AFDB dark clusters (Figure 7). In total, screening of 1,334,703 representatives across these three collections retained 70,560 candidate transition metal-binding sites, of which MAHOMES-II classified 8,961 as catalytic and 61,599 as non-catalytic.

**Figure 6.**
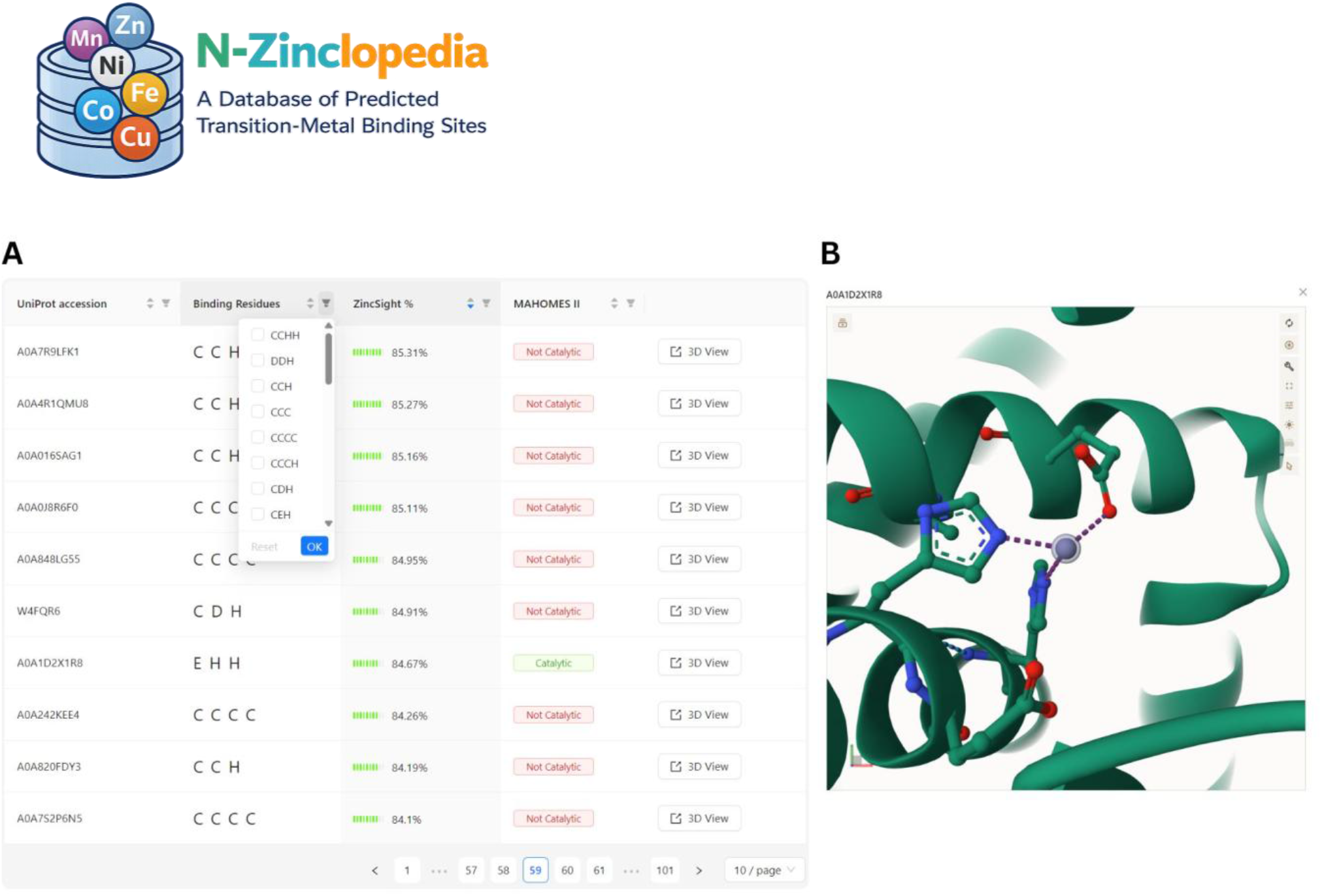
The N-Zinclopedia database interface. Left, a searchable and filterable results table lists the protein identifier, predicted coordinating residues, ZincSight score, and MAHOMES-II catalytic classification for each site. Right, an integrated Mol* 3D structure viewer displays the selected protein model with the predicted metal-binding site highlighted, enabling inspection of the local coordination environment.

**Figure 7.**
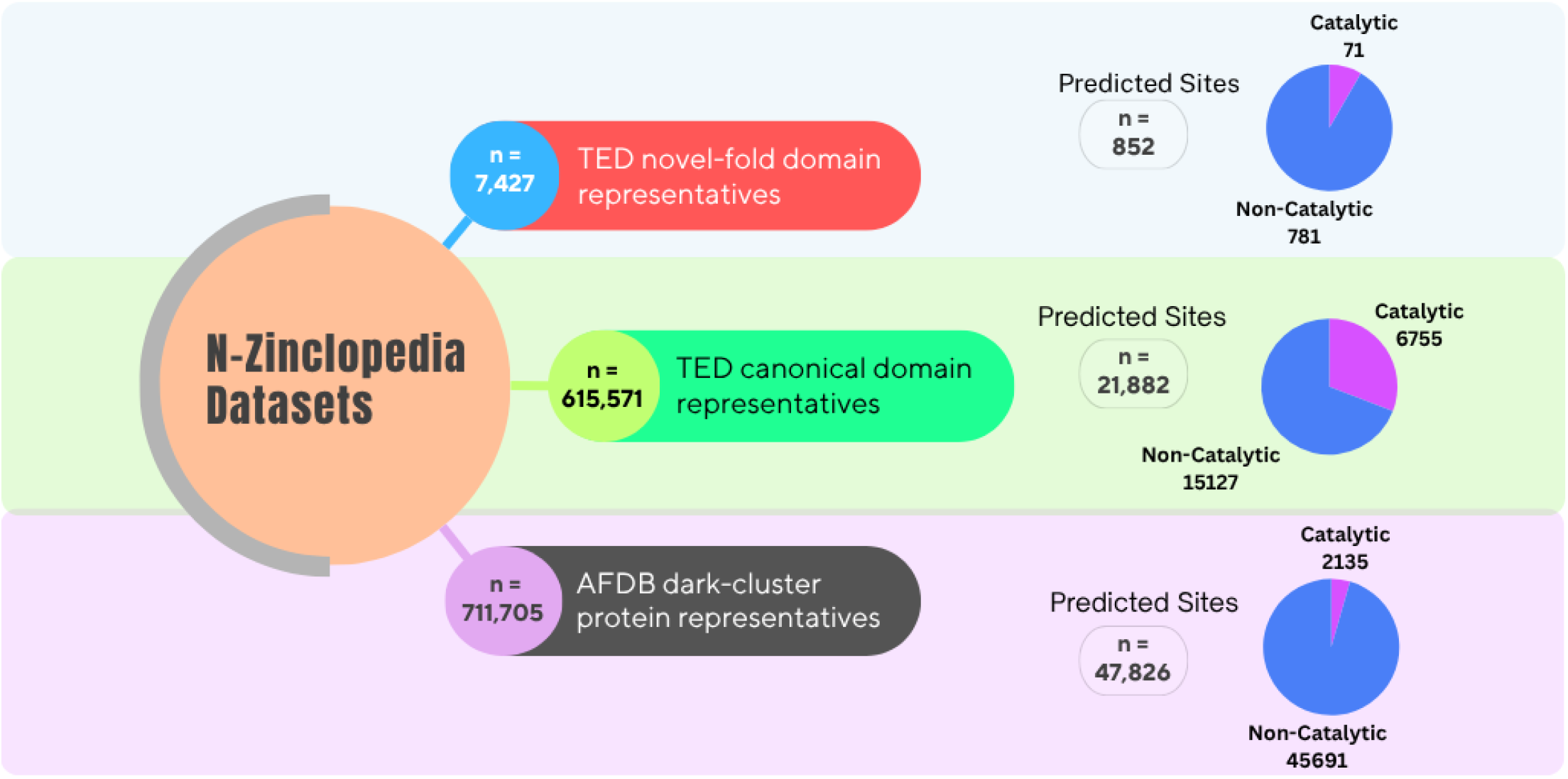
Overview of N-Zinclopedia input collections and predicted-site classification. N-Zinclopedia integrates three complementary structure collections comprising 1,334,703 representatives in total: 7,427 TED putative novel-fold domain representatives; 615,571 TED canonical-topology domain representatives, including 570,853 Rossmann-like domains (CATH 3.40.50), 32,271 TIM-barrel domains (CATH 3.20.20), and 12,447 seven-bladed β-propeller/methylamine dehydrogenase chain H-like domains (CATH 2.130.10); and 711,705 AFDB dark-clusters protein representatives. ZincSight was used to identify candidate transition-metal-binding sites in each collection, and MAHOMES-II subsequently classified the predicted sites as putatively catalytic (purple) or non-catalytic (blue). The TED putative novel-fold collection yielded 852 predicted sites, comprising 71 catalytic and 781 non-catalytic sites. The TED canonical-topology collection yielded 21,882 predicted sites, comprising 6,755 catalytic and 15,127 non-catalytic sites, whereas the AFDB dark-clusters collection yielded 47,826 predicted sites, comprising 2,135 catalytic and 45,691 non-catalytic sites. The values shown beside the dataset labels indicate the numbers of input domain or protein representatives, whereas the values beside the pie charts indicate the numbers of predicted sites.

## Discussion

N-Zinclopedia provides a database-scale atlas of predicted transition metal-binding sites across protein fold space. It serves as a discovery resource and a practical platform for downstream analysis. By combining ZincSight with MAHOMES-II, it not only identifies candidate metal-binding pockets but also prioritizes sites more likely to be catalytic, helping distinguish putative metalloenzymes from likely structural or regulatory metal-binding sites.

A central conclusion of this work is that unannotated candidate metalloenzymatic sites are broadly distributed across protein structure space, not only in putative novel folds but also in canonical topologies and annotation-poor proteins outside standard domain-resolved sets. This indicates that missing metal-related annotation is not restricted to rare structural novelty, but reflects a broader gap in functional mapping across diverse structural contexts.

The scientific value of N-Zinclopedia lies in enabling residue-level, cross-fold analysis of metal-binding pockets at scale. Because metal-dependent function is governed primarily by local structural and physicochemical features, including ligand composition, coordination geometry, inter-ligand spacing, pocket accessibility, and the surrounding microenvironment, a large atlas of candidate sites makes it possible to examine how these constraints recur across otherwise unrelated scaffolds. When these constraints are not satisfied, metal binding may fail or may not support productive catalysis, underscoring the importance of evaluating candidate sites in their full local structural context. In this sense, N-Zinclopedia is not only a catalog of predictions, but also a framework for studying general principles of metal-site architecture and metalloenzyme organization across fold space.

The atlas is also valuable in practical terms. By providing a large repertoire of candidate catalytic metal-binding pockets, it offers starting points for biochemical examination, comparative analysis, and metal-aware protein engineering. More broadly, it expands the searchable space of candidate scaffolds for reactions in which metal-centered chemistry may be advantageous or required, including Lewis-acid activation, metal-hydroxide chemistry, and redox-associated mechanisms. Although these applications remain prospective, N-Zinclopedia provides a practical framework for prioritizing candidates and pursuing them systematically.

Experimental characterization of A0A3W0S055 supported the N-Zinclopedia prediction of metal binding, with zinc being the predominant transition metal detected in the purified protein. Although N-Zinclopedia provided an annotation of two zinc ions per monomer of A0A3W0S055, we measured zinc-to-protein molar ratios of 0.5 before zinc loading and 0.91 after zinc loading and desalting. The measured ratios do not establish whether one or both predicted sites contribute to zinc binding. These experimental results demonstrate the utility of N-Zinclopedia, for guiding metalloprotein discovery, while also revealing a discrepancy between the predicted and experimentally measured zinc-to-protein ratios under the experimental conditions examined. It remains unclear whether one or both of the predicted sites contain zinc.

N-Zinclopedia should be interpreted as a hypothesis-generating resource rather than a definitive annotation set. The current workflow relies heavily on predicted structures, uses AlphaFold3 only as a targeted follow-up step, and applies MAHOMES-II to infer catalytic propensity from local site properties rather than direct biochemical activity. In addition, predicted site geometry alone does not establish physiological metal identity, occupancy, or function. It is also noteworthy that, ZincSight does not reliably distinguish zinc from other transition metals with similar coordination chemistry. Accordingly, entries in N-Zinclopedia are best interpreted as candidate transition metal-binding sites rather than definitive zinc-specific assignments, and the physiological metal identity of each site remains to be established experimentally. Experimental validation will therefore be essential for determining which predicted sites correspond to *bona fide* metalloenzymes.

Overall, N-Zinclopedia provides more than a collection of predicted sites: it offers a scalable way to organize, compare, and prioritize candidate metal-binding pockets across diverse structural contexts. Its main strength lies in linking atlas-scale coverage with residue-level functional hypotheses, thereby creating a resource that is valuable both for general analysis of biological metal utilization and for practical identification of experimentally tractable metalloenzyme candidates. As an open-source and extensible platform, it can be readily expanded to additional fold classes, annotation-poor protein collections, and other biologically relevant metals as new structural data accumulate, enabling the community to broaden coverage systematically over time.

## Methods

### TED novel-domain screening and candidate-site selection

A total of 7,427 representative TED novel-fold domains were screened with ZincSight to identify candidate zinc-binding sites. Scoring thresholds were established using the F₁ and recall-oriented Fβ (β = 2) calibration procedure described in the original ZincSight study (18). Because TED domains are largely derived from predicted structures, local side-chain inaccuracies can compromise metal-site detection, particularly for histidine residues whose imidazole orientations may be mis-modeled. To address this while avoiding overinterpretation of rotamer refinement alone, each candidate site was evaluated in two complementary modes.

In the first mode, sites were scored directly on the input structure without rotamer adjustment, and only those with a ZincSight probability of p ≥ 0.2 were retained. This baseline gate ensured that the modeled local geometry already exhibited an intrinsic predisposition for zinc coordination. In the second mode, histidine imidazole rotamer sampling was enabled to recover zinc-compatible geometries obscured by local side-chain inaccuracies; under this refinement mode, sites were required to satisfy a stricter threshold of p_rot_ ≥ 0.4. Only sites passing both gates were retained. This two-step strategy was designed to preserve a conservative baseline while allowing recovery of plausible metal-binding pockets from predicted structures. Requiring agreement between the unrefined and refined evaluations also reduced refinement-driven score inflation, in which a favorable score arises only after an expanded conformational search rather than from the original modeled pocket geometry. This dual-gate strategy therefore balances sensitivity for mis-modeled residues with a high specificity requirement, ensuring that the final candidate list is not dominated by artifacts of the rotamer optimization process. The same dual-gate ZincSight retention criteria were applied to the TED canonical-topology and AFDB dark-cluster collections.

### Catalytic-site prioritization with MAHOMES-II

Candidate zinc-binding sites were next evaluated with MAHOMES-II, which predicts catalytic propensity by analyzing features such as pocket depth, solvent accessibility, and the electrostatic environment of the coordination sphere. Sites with p_(MAHOMES)_ ≥ 0.5 were classified as putatively catalytic and were carried forward for subsequent novelty and validation analyses.

### Structural novelty assessment

To distinguish potentially unannotated metalloenzymes from sites readily explained by known structural homologues, putatively catalytic candidates were filtered by Foldseek structural comparison against annotated proteins. These searches covered all databases available through the Foldseek web server as of June 2026. Candidates with Foldseek structural neighbors assigned to previously characterized metalloenzymatic functions and match probability > 0.5 were removed. Foldseek was used as the principal novelty filter because structural comparison is substantially more sensitive than sequence-only similarity searches (e.g., BLAST, MMseqs2, HMMER) for detecting remote homology, while remaining computationally practical at this scale (28, 43). Candidates that remained unexplained after Foldseek filtering were therefore prioritized as stronger candidates for previously unannotated metalloenzymatic architectures.

### AlphaFold3 complex modeling and stoichiometry analysis

AlphaFold3 was not used in the initial high-throughput screening workflow, but only as a targeted follow-up step for candidates surviving Foldseek filtering. These candidates were modeled as protein–zinc complexes with AlphaFold3; zinc being placeholder for any transition metal. Confidence in the modeled metal placement was assessed using the local pLDDT of the zinc ion, and only sites with zinc-ion pLDDT ≥ 90 were retained; 12 sites falling below this threshold were excluded. For selected candidates in which the initial zinc-ion confidence was reduced, additional AlphaFold3 models were generated under alternative zinc stoichiometries to test whether local confidence improved with a two-zinc input. In these cases, ZincSight was also re-applied to the corresponding metal-free and metal-containing AlphaFold3 models (after metal removal). The idea was to assess whether the predicted pocket geometry was preserved across different metal-occupancy states, or whether a second site was only recoverable after initial metal binding.

### Evolutionary conservation analysis

Per amino acid site conservation was evaluated with the ConSurf pipeline, which automatically identifies homologues optimized for robust conservation profiling across diverse taxa. Homologues were collected using an E-value threshold of 1 × 10⁻⁴, a maximum pairwise sequence identity of 95%, and a minimum identity of 10%, aiming to retain broad evolutionary sampling while limiting overrepresentation of near-identical sequences. Predicted coordinating residues were examined for ConSurf conservation scores and category assignments indicative of functional constraint (Table 1).

### Functional annotation of representative candidates

For selected high-confidence candidates, additional tools were applied to supplement structural inference. Gene Ontology molecular function and biological process terms were predicted from protein structure using DeepFRI (37). Enzyme class membership and candidate EC numbers were predicted with EZPred (38). Predicted functional protein–protein associations were retrieved from STRING (35) to provide network-level context for possible pathway membership.

### Cloning and Overexpression

DNA was subcloned into the pET-11a vector to enable overexpression of target proteins using isopropyl β-D-1-thiogalactopyranoside (IPTG). Plasmids containing the protein-coding sequences were transformed into *E. coli* strain BL21 (DE3) for IPTG-inducible overexpression. Small-scale (25 mL) cultures of Luria-Bertani (LB) broth containing 100 µg/mL carbenicillin were inoculated and grown overnight at 37° C. These cultures were used to inoculate large-scale (1 L) cultures of Terrific Broth (TB) containing 100 µg/mL carbenicillin and grown in a shaking incubator at 37° C/300 rotations per minute (RPM). Large-scale cultures were grown to an optical density (A_600_) of ∼1.3 - 1.6 before being rapidly cooled to 20° C in an ice-water bath. Cultures were induced with 100 µM IPTG and allowed to express for 20 h in a shaking incubator at 20° C/300 RPM. Post-expression cultures were cooled to 4° C in an ice-water bath and harvested by centrifugation at 4,000 x g. Pellets were frozen and stored at -80° C for subsequent purification.

### Protein Purification

Cell pellets were thawed and thoroughly resuspended in lysis buffer (50 mM Tris-HCl pH 8.0 at 4° C) to a final volume of 45 mL. DNase I was added to ≥5 µg/mL to cleave genomic DNA. Resulting cell suspensions were placed in a rigid ice-water bath and sonicated using a QSonica Q500 Sonicator (2:00 duration, 40% amplitude, 2s/8s duty cycle). Lysate was clarified by centrifugation at 48,384 x g at 4° C. The supernatant was carefully extracted for further purification.

Ammonium sulfate precipitation was used for the first level of purification (Burgess 2009) of A0A3W0S055. Solid high-purity ammonium sulfate (≥99%, Sigma-Aldrich) used for protein purification was crushed into a fine powder using a mortar and pestle to promote rapid dissolution of the powder and prevent unintended precipitation of the target. Clarified lysate was adjusted to a final volume of 50 mL with lysis buffer and placed on ice in a stirred beaker. Solid ammonium sulfate was slowly added to 50% saturation at 0° C (14.55 g), and the solution was allowed to equilibrate (stirred, on ice) for 30 min. After equilibration, insoluble material was removed by centrifugation at 10,000 x g for 10 min at 4° C. The supernatant was removed to a new beaker, and ammonium sulfate was added to a final saturation of 60% at 0° C (3.4981 g). Ammonium sulfate-precipitated pellets were redissolved in ∼10 mL of lysis buffer; the presence of the protein in the 50%-60% ammonium sulfate was confirmed by SDS-PAGE.

The redissolved protein solution was dialyzed twice against 4 L lysis buffer to remove any residual ammonium sulfate prior to performing anion exchange chromatography – once for 3 h at room temperature, and again overnight. Following dialysis, the solution was centrifuged at 14,000 x g for 10 min to remove insoluble material. The pellet was dissolved in 8M urea, and both pellet and supernatant were analyzed by SDS-PAGE to confirm the presence of protein in the soluble fraction. The soluble fraction was applied to a 5 mL HiTrapQ HP column (Cytiva) and purified in 20 mM Tris-HCl pH 8.0 using a linear NaCl gradient from 0 – 1 M on an AKTA Pure 25 chromatography system (Cytiva) at room temperature. Fractions from the most prominent peak were pooled, concentrated to a volume of ∼500 µL using an Amicon ultracentrifugal filter (Millipore Sigma), and further purified by size-exclusion chromatography using a Superdex 200 Increase 10/300 GL column (Cytiva) using 20 mM Tris-HCl pH 8.0 150 mM NaCl as the running buffer at a flow rate of 1.0 mL/min at room temperature. The protein concentration and yield were determined by A_280_ using the theoretical molar extinction coefficient provided by ExPASy ProtParam (32,420 M^-1^ cm^-1^).

For creating zinc-soaked A0A3W0S055, protein was soaked in excess zinc, 1 mM ZnCl2 and 60 µM protein, for one hour and then desalted into 20 mM Tris-HCl pH 8.0 150 mM NaCl using a 5 mL HiTrap desalt column (Sephadex G-25, Cytiva) (5.0 mL/min) at room temperature.

### Mass Spectrometry

The intact mass of A0A3W0S055 was measured by MALDI (matrix-assisted laser desorption/ionization) mass spectrometry using a Bruker Autoflex Max using a matrix of 10 mg/mL α-cyano-4-hydroxycinnamic acid (HCCA) in 50% (v/v) acetonitrile.

### Inductively Coupled Plasma Optical Emission Spectroscopy (ICP-OES)

A0A3W0S055 in 20 mM Tris-HCl pH 8.0 150 mM NaCl was transferred to the digestion vessel. Then, 5 mL of nitric acid (70%), and 2 mL of 18 MΩ Millipore water were added. The samples were digested in the CEM MARS6 microwave digestion system using the following parameters: Power 700-1800 W, Temperature 240 o C, Ramp Time 30 mins, Hold Time 15 mins

After digestion, the samples solutions were diluted to a final volume of 25 mL. A stock standard of 5 ppm was prepared in 2% nitric acid. The Agilent ADS 2 was used to generate a 6-point calibration curve ranging from 0.05 to 5 ppm. Samples and standards were analyzed on an Agilent 5900 ICP-OES instrument with the following operating parameters and a high purge rate:Plasma (Argon) gas flow 14 L/min, Auxiliary gas flow 1 L/min, Nebulizer gas flow 0.80 L/min, RF Power 1200 W

Several emission lines were measured for the element of interest. The line that presented the highest precision and recovery was used for quantification (206.200 nm). A quality control standard was run after the samples, which was within 10% of the expected concentration.

For the semiquantitative analysis, a 5 ppm multielement standard and blank were analyzed using at least three wavelengths for every element.

## Acknowledgements

This work was supported by Israel Science Foundation (ISF) grant 1764/21, and by the Minerva Foundation. Nir Ben-Tal’s research is supported by the Abraham E. Kazan Chair in Structural Biology, Tel Aviv University. G.M. and G.A. were supported in part by a fellowship from the Edmond J. Safra Center for Bioinformatics at Tel-Aviv University.

We gratefully thank Ashley Blystone at the Microanalysis Laboratory, University of Illinois for ICP-OES experimentation and NSF award 2226804 for funding.

